# Widespread drastic reduction of brain myelin content upon prolonged endurance exercise

**DOI:** 10.1101/2023.10.10.561303

**Authors:** Pedro Ramos-Cabrer, Alberto Cabrera-Zubizarreta, Daniel Padró, Mario Matute-González, Alfredo Rodríguez-Antigüedad, Carlos Matute

## Abstract

Recent evidence suggests that myelin lipids may act as glial energy reserves when glucose is lacking, a hypothesis yet to be solidly proven. Hereby, we examined the effects of running a marathon on the myelin content by MRI. Our findings show that marathon runners undergo widespread robust myelin decrease at completion of the effort. This reduction involves white and gray matter, and includes primary motor and sensory cortical areas and pathways, as well as the entire corpus callosum and internal capsule. Notably, myelin levels partially recover within two weeks after the marathon. These results reveal that myelin use and replenishment is an unprecedented form of metabolic plasticity aimed to maintain brain function during extreme conditions.

**One-Sentence Summary:** Brain myelin usage during strenuous exercise and recovery thereafter

---

Prolonged endurance exercise mobilizes energy stores throughout the body to meet energy demands. Carbohydrates are the primary fuel source which is provided by glycogen breakdown in muscles and liver. As glycogen stores become depleted, the body starts to rely more on stored fat as an energy source. Triglycerides are broken down into fatty acids, which are released from adipose tissue to produce energy by beta-oxidation. Ultimately and if needed, the body may break down muscle protein to use as energy by gluconeogenesis. During strenuous exercise, all these energy sources may be utilized simultaneously or in varying proportions depending on factors like exercise intensity, duration, individual fitness level and training routines.

Marathon runners are not an exception to these rules, and primarily rely on carbohydrates as the main energy source during a marathon race (*1*). As the race progresses and glycogen stores are depleted, marathon runners use fat as an energy source (*1,2*). Fat is a more abundant energy source in the body compared to carbohydrates, and it can provide a sustained source of energy for prolonged endurance exercise. The utilization of fat as an energy source increases as the race goes on and glycogen stores become depleted. In the rat brain, astrocytic glycogen-derived lactate contributes to energy expenditure during exhaustive exercise (*3,4*).

Myelin surrounds and enwraps axons in the central and peripheral nervous systems providing electrical insulation and metabolic support to axons. The primary components of myelin are lipids, which make up 70-80% of it, whereas proteins compact and stabilize its multi-layered structure. While myelin lipids have been hypothesized to serve as glial energy reserves, in a mutant mouse model of oligodendroglial glucose deprivation (*5*), here we further hypothesize that myelin lipids may contribute to brain activity as body fat does to fuel muscle energy demands.

To test our hypothesis, we evaluated myelin levels at full brain scale, by Multicomponent Relaxometry Magnetic Resonance Imaging in city and mountain marathon runners. To that end, we acquired multi-echo T2 weighted MRI sequences, postprocessed with the DECAES algorithm for Multicomponent Relaxometry analysis (*6*) thus providing 3D parametric maps of Myelin Water Fraction (MWF). MWF has been purported to measure pools of water molecules within myelin lamellae, as a proxy, and it is therefore a biomarker for myelination (*6–8*) with high sensitivity to detect subtle variations in myelin content (*9*). In this manner, we estimated water trapped between the lipid bilayers of the myelin sheath around axons and excluded luminal water that accounts for intra- and extracellular water (fig. S1).

## Running a marathon robustly lowers brain myelin content

MRI based MWF maps were constructed for the 4 subjects within 48 h before running a marathon. All individuals showed similar distribution of myelin across the brain, with naturally occurring interindividual variabilities (Fig. 1A). Maps of luminal water fraction (LWF) and myelin fraction T2 times and luminal fraction T2 times were also obtained in the fitting process (figs. S2 and S3). High resolution 3D T1 weighted MRI images of all subjects showed no significant interindividual differences for the different imaging sessions (fig. S4). Absence of large differences among subjects allowed us to build up averaged (mean, with n=4) MWF maps of the brain, as presented in Figure 1B, with MWF values in the range 0 - 0.25 for gray matter regions and 0.2 - 0.55 for large white matter tracks, which are in line with MWF values found in healthy subjects on a wide range of ages (*10,11*).

**Figure 1.**
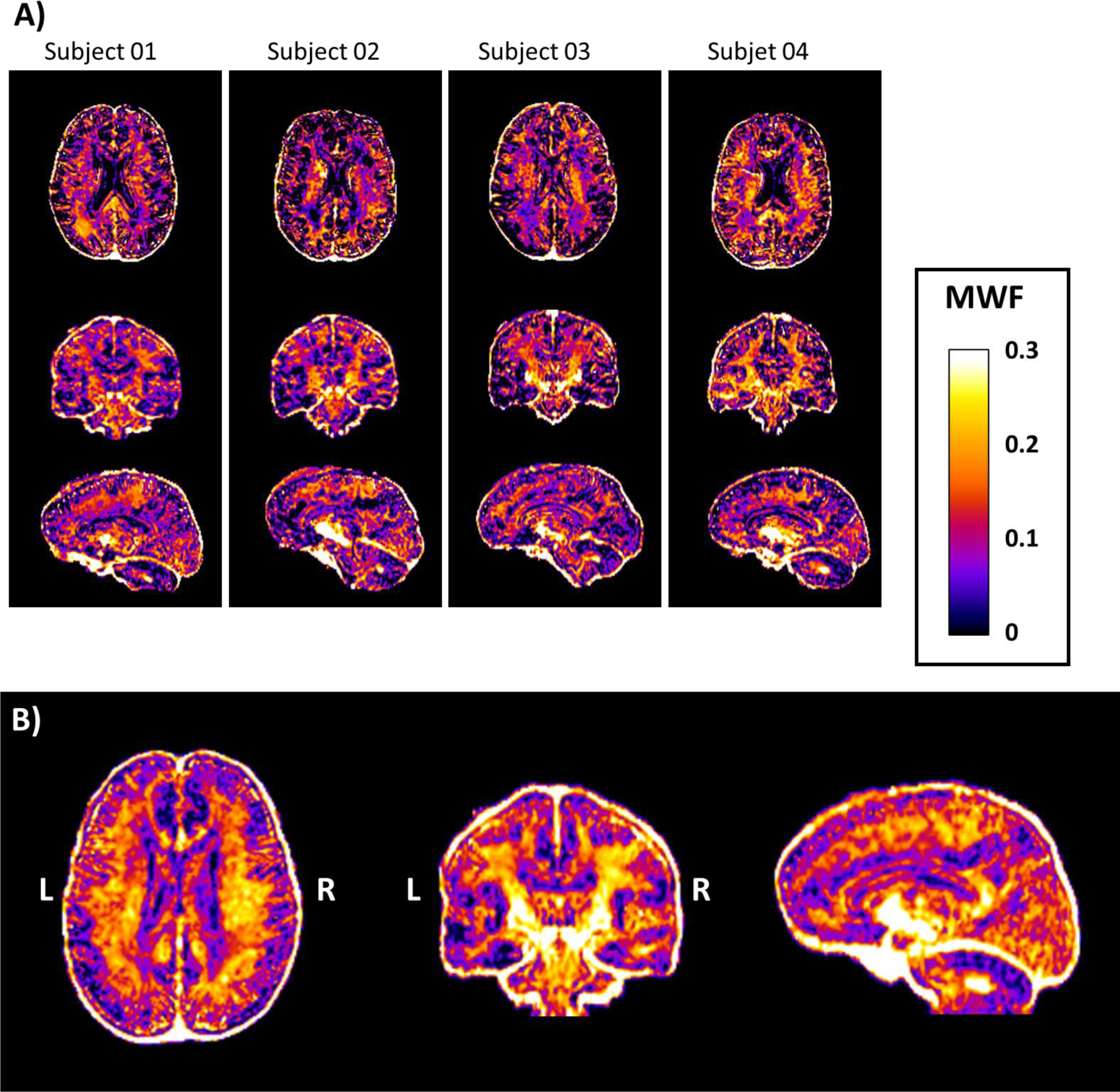
Myelin Water Fraction (MWF) parametric maps from subjects before the marathon challenge. A) Pre-exercise MWF maps for the 4 subjects scanned in this study. From top to bottom, arbitrarily chosen axial, coronal and sagittal planes. B) Average maps (mean of n=4 subjects) obtained after ANTs Syn registration of all images to a common spatial frame.

Similar to Figure 1B, averaged maps of MWF values were constructed for the second (n=4 individuals) and third (n=2 individuals) imaging sessions, corresponding to post-exercise (24-48 h after the marathon) and after recovery (2 weeks later) statuses. Coronal sections of three imaging sessions are presented in Figure 2A, showing a clear post-exercise reduction of WMF in the motor descending pathways, among other regions. Total brain, CSF, gray matter, white matter, deep brain, brainstem and cerebellum volume were not altered intra-individually across imaging sections (table S1).

**Figure 2.**
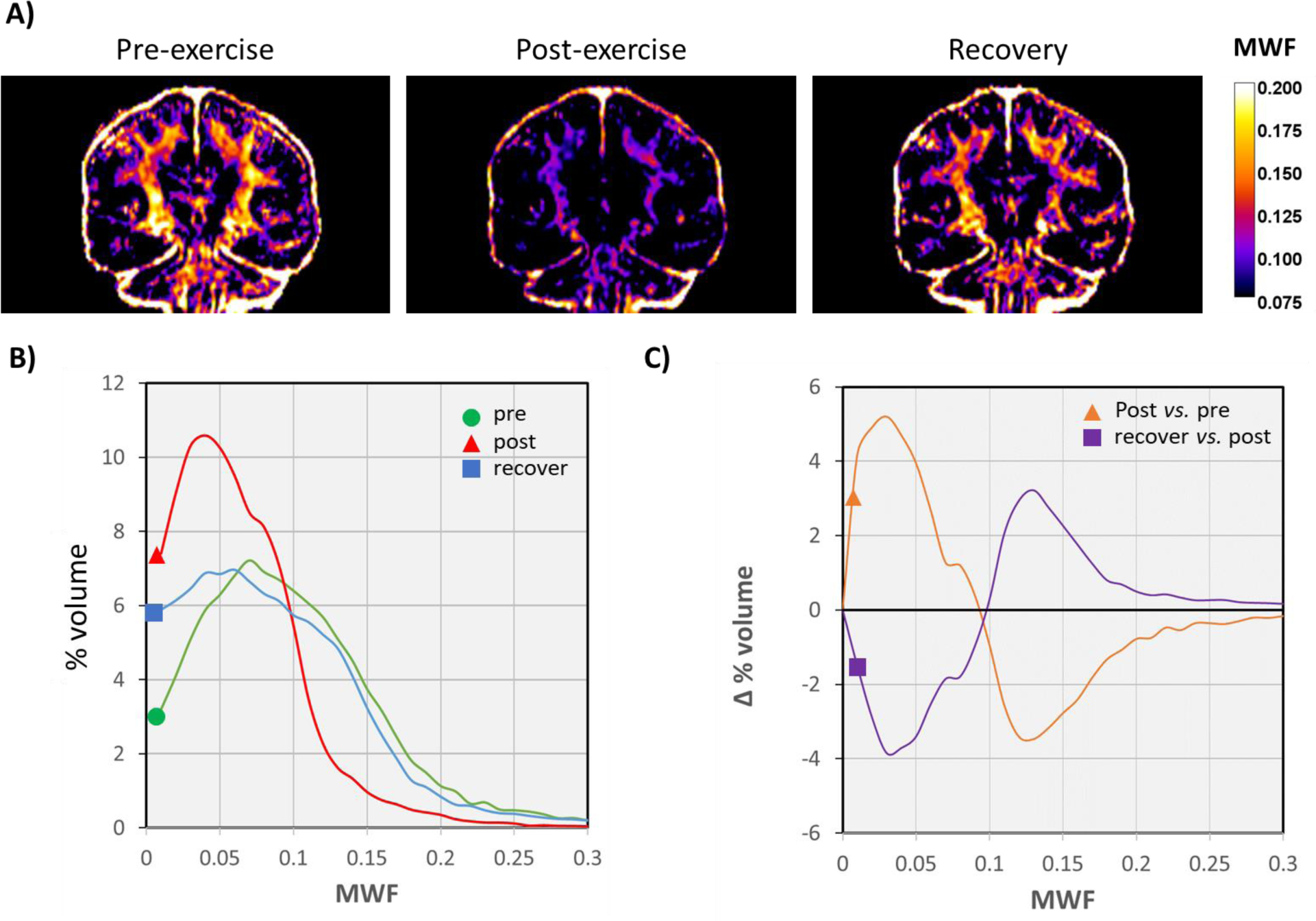
Myelin water fraction (MWF) in motor descending pathways. **A)** Coronal sections of MWF maps (mean of all individuals) pre-, post-exercise and after two weeks of recovery, showing areas with MWF values higher than 0.075. **B)** Curves plots showing percentage of volume presenting different MWF values in both hemispheres pooled together, in a slab of 5 coronal sections (centered at images presented in A), pre-(green plot, circle), post-exercise (red plot, triangle) and after recovery (blue curve, square). Each plot corresponds to the mean of all subject individual histograms for each imaging session. **C)** Difference in % of the slab volume, comparing imaging session Post-vs. Pre-exercise (orange curve, triangle) and recovery vs. post-exercise (purple curve, square).

## Myelin levels swiftly rise during post-marathon recovery

In order to quantify the observed myelin reduction and subsequent status of MWF after prolonged endurance exercise, two different strategies were followed.

In the first place, we built curves accounting for regions with different MWF values in both hemispheres (Fig. 2B) from 5 consecutive coronal sections containing the pyramidal tract arising from motor cortex towards brainstem. Plots (expressed as % of volume) were constructed for the inter-subject averaged MWF maps, at the three imaging sessions (pre-exercise: green plot; post-exercise: red plot; and after recovery: blue plot; Fig. 2B). Before exercise MWF values ranged from 0 to 0.3 with an area under the curve of 98.85% and peaking at MWF=0.07 with a value of volume of 7.23%. After running the marathon, myelin content was robustly reduced. Thus, MWF curve shifted to the left, presented an area under the curve of 99.84% and peaked at MWF=0.04, with a volume of 10.59%. Comparison of both plots illustrates how regions with MWF higher than 0.1 drops after exercise, increasing the areas with MWF<0.1 (Fig. 2B). Conversely, after two weeks of recovery MWF curve shifted back towards higher values presenting an area under the curve of 98.41% and peaking at MWF=0.06, with a value of 6.96%. Despite initial pre-exercise myelin content values not being reached, these results indicate a swift and nearly complete myelin rebuilding in the pyramidal tract towards the initial status levels after two weeks of resting. Analysis of the data showed a marked decrease of areas with higher MWF values along with a rise in areas with lower MWF values after exercise (orange curve; Fig. 2C). The opposite distribution was observed after two weeks of recovery and therefore, demonstrated a clear increase in regional volume with higher MWF (purple curve; Fig. 2C) post-recovery.

## Global widespread brain myelin reduction and recovery after strenuous exercise

Next, to get a comprehensive view of myelin content changes we performed an analysis of MWF values after segmentation of the entire brain into different regions with the help of an atlas-aided 3D morphometric analysis. Thus, in this second approach, we segmented the brain in 50 white matter regions, using the JHU brain atlas of white matter tracks (*12*) and 56 gray matter regions, using the LPBA40 collection of the SRI24 brain atlas (*13*). Mean MWF values were recorded for these 106 brain regions for each subject at each imaging session (Table 1 and table S2; Fig. 3).

**Figure 3.**
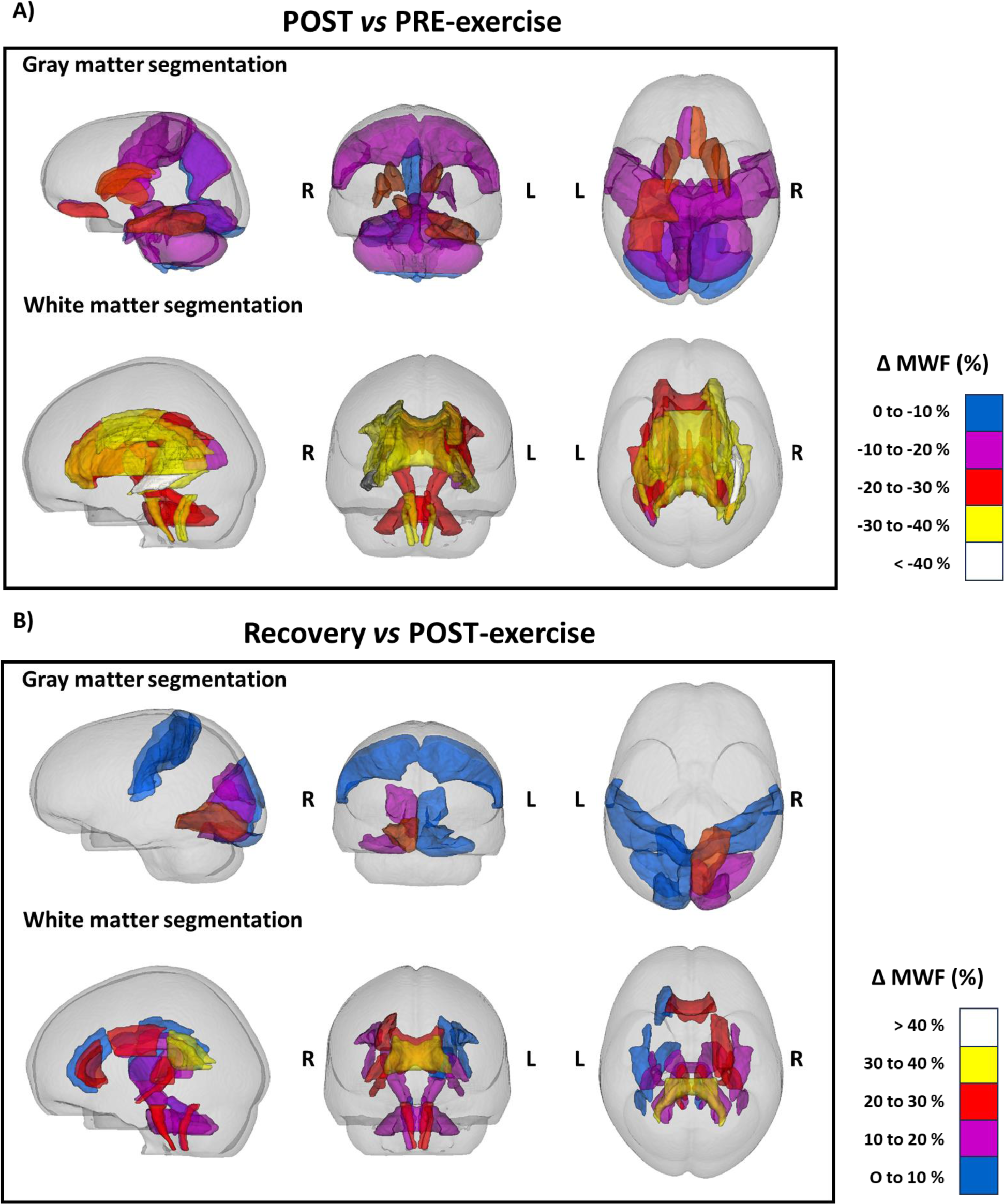
Color encoded representations in 3D of regions of the brain that show significant reduction of Myelin Water Fraction post-exercise (**A**), and increase after recovery (**B**). Segmented white matter regions are displayed using the JHU atlas while segmented gray matter regions are displayed using the LPBA40 set of the SRI24 atlas.

**Table 1.** Changes of MWF from pre-to post-exercise. Only regions with WMF values higher than 0.07 in either hemisphere at pre-exercise stage (n=4) and with changes higher than 5% are included (see table S2 for a complete data set). L, left; R, right; SD, standard deviation. Areas determined according to maps in refs. *12* and *13*.

| Gray matter |  |  |  |  |  |
| --- | --- | --- | --- | --- | --- |
| Region | MWF pre-run | | MWF post-run | | ΔMWF (%) |
|  | mean | SD | mean | SD |  |
| L precentral_gyrus | 0.107 | 0.016 | 0.089 | 0.008 | -15 |
| R precentral_gyrus | 0.097 | 0.014 | 0.083 | 0.008 | -14 |
| L gyrus_rectus | 0.082 | 0.027 | 0.065 | 0.025 | -21 |
| R gyrus_rectus | 0.070 | 0.035 | 0.059 | 0.025 | -12 |
| L postcentral_gyrus | 0.103 | 0.011 | 0.085 | 0.007 | -17 |
| R postcentral_gyrus | 0.096 | 0.018 | 0.081 | 0.016 | -15 |
| L precuneus | 0.076 | 0.013 | 0.066 | 0.015 | -10 |
| R precuneus | 0.088 | 0.027 | 0.073 | 0.015 | -12 |
| L inferior_occipital_gyrus | 0.117 | 0.020 | 0.107 | 0.026 | -7 |
| R inferior_occipital_gyrus | 0.174 | 0.029 | 0.157 | 0.025 | -9 |
| L parahippocampal_gyrus | 0.089 | 0.012 | 0.078 | 0.016 | -13 |
| R parahippocampal_gyrus | 0.116 | 0.017 | 0.093 | 0.021 | -21 |
| L lingual_gyrus | 0.137 | 0.036 | 0.121 | 0.022 | -10 |
| R lingual_gyrus | 0.143 | 0.026 | 0.118 | 0.023 | -17 |
| L fusiform_gyrus | 0.062 | 0.011 | 0.052 | 0.013 | -16 |
| R fusiform_gyrus | 0.091 | 0.025 | 0.066 | 0.026 | -28 |
| L caudate | 0.098 | 0.023 | 0.070 | 0.022 | -25 |
| R caudate | 0.089 | 0.029 | 0.068 | 0.022 | -21 |

| **L putamen** | 0.200 | 0.043 | 0.155 | 0.025 | -22 |
| --- | --- | --- | --- | --- | --- |
| **R putamen** | 0.205 | 0.031 | 0.169 | 0.021 | -17 |
| **cerebellum** | 0.073 | 0.011 | 0.059 | 0.018 | -20 |
| **brainstem** | 0.132 | 0.012 | 0.108 | 0.020 | -18 |
| **White matter** |  |  |  |  |  |
| **Region** | **MWF pre-run** |  | **MWF post-run** |  | **ΔMWF (%)** |
|  | **mean** | **SDEV** | **mean** | **SDEV** |  |
| **Middle cerebellar peduncle** | 0.109 | 0.019 | 0.079 | 0.026 | -29 |
| **Pontine crossing tract** | 0.134 | 0.013 | 0.106 | 0.056 | -23 |
| **Genu of corpus callosum** | 0.105 | 0.015 | 0.077 | 0.028 | -28 |
| **Body of corpus callosum** | 0.080 | 0.008 | 0.055 | 0.016 | -31 |
| **Splenium of corpus callosum** | 0.123 | 0.033 | 0.079 | 0.029 | -37 |
| **Fornix** | 0.058 | 0.020 | 0.046 | 0.023 | -24 |
| **R Corticospinal tract** | 0.133 | 0.019 | 0.095 | 0.041 | -29 |
| **L Corticospinal tract** | 0.141 | 0.007 | 0.097 | 0.042 | -32 |
| **R Medial lemniscus** | 0.105 | 0.014 | 0.086 | 0.035 | -20 |
| **L Medial lemniscus** | 0.118 | 0.026 | 0.094 | 0.045 | -24 |
| **R Inferior cerebellar peduncle** | 0.144 | 0.023 | 0.104 | 0.046 | -36 |
| **L Inferior cerebellar peduncle** | 0.134 | 0.016 | 0.098 | 0.040 | -33 |
| **R Superior cerebellar peduncle** | 0.248 | 0.047 | 0.186 | 0.055 | -29 |
| **L Superior cerebellar peduncle** | 0.244 | 0.044 | 0.182 | 0.044 | -29 |
| **R Cerebral peduncle** | 0.162 | 0.043 | 0.112 | 0.019 | -23 |
| **L Cerebral peduncle** | 0.162 | 0.038 | 0.117 | 0.011 | -24 |
| **R Anterior limb of int. capsule** | 0.194 | 0.040 | 0.133 | 0.040 | -28 |
| **L Anterior limb of int. capsule** | 0.205 | 0.023 | 0.142 | 0.034 | -25 |
| **R Posterior limb of int. capsule** | 0.131 | 0.042 | 0.081 | 0.021 | -30 |
| **L Posterior limb of int. capsule** | 0.143 | 0.028 | 0.090 | 0.028 | -31 |
| **R Retrolenticular part of int. capsule** | 0.100 | 0.007 | 0.072 | 0.037 | -36 |
| **L Retrolenticular part of int. capsule** | 0.081 | 0.021 | 0.052 | 0.020 | -38 |
| **R Anterior corona radiata** | 0.129 | 0.012 | 0.087 | 0.040 | -29 |
| **L Anterior corona radiata** | 0.111 | 0.027 | 0.079 | 0.039 | -37 |
| **R Superior corona radiata** | 0.087 | 0.018 | 0.066 | 0.019 | -32 |
| **L Superior corona radiata** | 0.100 | 0.018 | 0.072 | 0.030 | -31 |
| **R Posterior corona radiata** | 0.081 | 0.025 | 0.066 | 0.035 | -25 |
| **L Posterior corona radiata** | 0.097 | 0.010 | 0.060 | 0.020 | -30 |
| **R Posterior thalamic radiation** | 0.093 | 0.046 | 0.060 | 0.038 | -18 |
| **L Posterior thalamic radiation** | 0.093 | 0.026 | 0.055 | 0.024 | -39 |
| **R Sagittal stratum** | 0.104 | 0.023 | 0.076 | 0.003 | -35 |
| **L Sagittal stratum** | 0.100 | 0.016 | 0.061 | 0.016 | -42 |
| **R External capsule** | 0.120 | 0.046 | 0.073 | 0.023 | -25 |
| **L External capsule** | 0.124 | 0.031 | 0.076 | 0.016 | -38 |
| **R Fornix (cres) / Stria terminalis** | 0.123 | 0.013 | 0.095 | 0.041 | -37 |
| **L Fornix (cres) / Stria terminalis** | 0.114 | 0.018 | 0.080 | 0.039 | -37 |
| **R Sup. longitudinal fasciculus** | 0.088 | 0.013 | 0.053 | 0.025 | -24 |
| **L Sup. longitudinal fasciculus** | 0.081 | 0.027 | 0.058 | 0.023 | -32 |
| **R Superior fronto-occipital fasciculus** | 0.073 | 0.014 | 0.060 | 0.027 | -39 |
| **L Superior fronto-occipital fasciculus** | 0.071 | 0.011 | 0.041 | 0.025 | -25 |

As changes in low myelin-containing areas can be subject to errors (see complete data set in table S2), we focused our attention on the regions of higher MWF values, as variations in this parameter are reflected better in more densely myelinated regions (*14*). Thus, areas with MWF values < 0.075 at the pre-run stage in all subjects under study are not included in Table 1, which summarizes quantitative regional MWF reduction in gray and white matter, while Table 2 illustrates 2-week post-marathon recovery.

**Table 2.** Changes of MWF from post-exercise to 2-week recovery after the marathon. Only regions with WMF values higher than 0.075 in either hemisphere at pre-exercise stage (n=2) are included (see table S2 for a complete data set). L, left; R, right; SD, standard deviation. Areas determined according to maps in refs. *12* and *13*.

| **Gray matter** |  |  |  |  |  |
| --- | --- | --- | --- | --- | --- |
| **Region** | **MWF post-run** |  | **MWF 2-week recovery** |  | **ΔMWF (%)** |
|  | **mean** | **SD** | **mean** | **SD** |  |
| **L postcentral_gyrus** | 0.087 | 0.005 | 0.090 | 0.004 | 4 |
| **R postcentral_gyrus** | 0.080 | 0.008 | 0.088 | 0.009 | 9 |
| **L superior_occipital_gyrus** | 0.083 | 0.006 | 0.097 | 0.006 | 15 |
| **R superior_occipital_gyrus** | 0.084 | 0.013 | 0.084 | 0.007 | 0 |
| **L inferior_occipital_gyrus** | 0.086 | 0.023 | 0.099 | 0.033 | 13 |
| **R inferior_occipital_gyrus** | 0.153 | 0.038 | 0.152 | 0.021 | 0 |
| **L cuneus** | 0.103 | 0.029 | 0.128 | 0.036 | 20 |
| **R cuneus** | 0.145 | 0.008 | 0.137 | 0.003 | -6 |
| **L lingual_gyrus** | 0.093 | 0.022 | 0.116 | 0.037 | 21 |
| **R lingual_gyrus** | 0.114 | 0.029 | 0.118 | 0.040 | 3 |
| **White matter** |  |  |  |  |  |
| **Region** | **MWF post-run** |  | **MWF 2-week recovery** |  | **ΔMWF (%)** |
|  | **mean** | **SD** | **mean** | **SD** |  |
| **Middle cerebellar peduncle** | 0.091 | 0.017 | 0.106 | 0.011 | 16 |
| **Genu of corpus callosum** | 0.079 | 0.040 | 0.097 | 0.049 | 23 |
| **Splenium of corpus callosum** | 0.083 | 0.045 | 0.109 | 0.031 | 31 |
| **R Corticospinal tract** | 0.113 | 0.001 | 0.139 | 0.006 | 24 |
| **L Corticospinal tract** | 0.104 | 0.011 | 0.126 | 0.001 | 21 |
| **R Medial lemniscus** | 0.095 | 0.034 | 0.104 | 0.006 | 10 |
| **L Medial lemniscus** | 0.100 | 0.034 | 0.110 | 0.005 | 10 |
| **R Inferior cerebellar peduncle** | 0.089 | 0.032 | 0.109 | 0.014 | 23 |
| **L Inferior cerebellar peduncle** | 0.088 | 0.019 | 0.103 | 0.001 | 17 |
| **R Superior cerebellar peduncle** | 0.125 | 0.041 | 0.141 | 0.028 | 13 |
| **L Superior cerebellar peduncle** | 0.116 | 0.035 | 0.120 | 0.001 | 4 |
| **R Cerebral peduncle** | 0.222 | 0.052 | 0.244 | 0.024 | 10 |
| **L Cerebral peduncle** | 0.211 | 0.038 | 0.248 | 0.017 | 18 |
| **R Posterior limb of internal capsule** | 0.162 | 0.023 | 0.169 | 0.015 | 4 |
| **L Posterior limb of internal capsule** | 0.168 | 0.012 | 0.189 | 0.004 | 12 |
| **R Retrolenticular part of int. capsule** | 0.089 | 0.021 | 0.100 | 0.018 | 11 |
| **L Retrolenticular part of int. capsule** | 0.105 | 0.016 | 0.131 | 0.007 | 25 |
| **R Anterior corona radiata** | 0.075 | 0.033 | 0.077 | 0.034 | 2 |
| **L Superior corona radiata** | 0.109 | 0.018 | 0.102 | 0.010 | 21 |
| **R Fornix (cres) / Stria terminalis** | 0.073 | 0.038 | 0.082 | 0.044 | 11 |
| **L Fornix (cres) / Stria terminalis** | 0.078 | 0.024 | 0.101 | 0.006 | 30 |
| **R Superior longitudinal fasciculus** | 0.114 | 0.044 | 0.114 | 0.046 | 0 |
| **L Superior longitudinal fasciculus** | 0.094 | 0.037 | 0.108 | 0.021 | 6 |

We observed consistent, region-variable MWF reductions (up to 28% in gray matter, and up to 42% in white matter) in many of the areas analyzed both in gray (12 out 56 areas) and in white matter (23 out of 50 areas; Fig. 3; Table 1). The amplitude of the changes was lower in gray matter than in white matter which likely reflects a lesser myelin content along with a reduced resolution to monitor MWF.

Gray matter changes after completion of exercise were prominent in cortical and subcortical brain regions involved in motor control including precentral gyrus, basal ganglia, brainstem and cerebellum. In addition, MWF lowered in postcentral, fusiform, precuneus, rectus, parahippocampal, lingual and inferior occipital and gyri. Thus, myelin reduction occurs bilaterally and with a similar amplitude in each hemisphere, in all brain lobes including motor, primary and secondary sensory gyri as well as gyri involved in higher level integration and cognition (Table 1). In turn, we observed a modest myelin increase in gray matter (up to 21%) after two weeks of post-marathon recovery, which was restricted to postcentral, occipital and lingual gyri (Table 2, table S2). Together, these findings indicate a long-lasting profound myelin remodeling in those areas, that may subtly modify functions associated with these gyri (*15, 16, 17*). It would be therefore important to evaluate whether this feature impacts on the neurophysiology and cognitive functions associated with those regions.

Strikingly, bilateral myelin loss in white matter was extensive and massive after completion of the marathon, and its extent was similar in both hemispheres. It involved all major tracts including the whole corpus callosum, fornix, internal capsule and corona radiata; corticospinal tracts, cerebral peduncles, all three cerebellar peduncles and medial lemnisci; and associative axonal tracts counting sagittal stratum (optic radiation and inferior longitudinal bundle), superior longitudinal fasciculi and superior fronto-occipital fasciculi (Fig. 2-3 and Table 1). Thus, MWF robustly diminished in major descending and ascending tracts, interhemispheric connections throughout the entire corpus callosum and ipsilateral association longitudinal tracts. Notably, high relative loss of myelin occurred in the superior longitudinal fasciculi, key ipsilateral connecting bundles interconnecting frontal, parietal, and medial temporal sites (*18*); and the fornix, the major output of the hippocampus and a key white matter tract of the limbic system participating in cognition and episodic memory recall (*19*). Importantly, white matter MWF values substantially increased (up to 31%) from the completion of the marathon to two weeks after the run in the corpus callosum, corticospinal tract, fornix/stria terminalis, internal capsule, and cerebellar peduncles, among other myelinated tracts (Fig. 3, Table 2 and table S2). Collectively, these findings provide compelling evidence that white matter tracts undergo a profound reduction of myelin content that it is restored later on after recovery from prolonged endurance exercise.

## Discussion

Our findings show that running a marathon reduces runners’ myelin content along the entire gray and white matter in the brain, in a region dependent manner and with a similar impact in both hemispheres. This myelin loss is partially restored thereafter as MWF values steadily increase at two weeks after completion of the effort. This widespread drastic reduction of myelin content upon prolonged exercise, and recovery after lowering physical activity, opens a novel view of myelin as an energy store ready to use when typically used brain nutrients are in short supply. We name this process metabolic myelin plasticity.

To our knowledge, myelin load has not been evaluated so far following strenuous prolonged exercise. Subtle reversible volume decreases in global gray matter, but not in white matter, in ultramarathon multistage runners have been reported earlier with no other signs of brain changes and damage, determined by MRI and evaluations for neurological complications (*20–23*). This reduction and subsequent recovery were paralleled by a similar loss in body weight and was gained back following a period of rest (*21*). Interestingly, EEG activity increases bilaterally in all brain lobes during graded exercise, correlates with its intensity, and returns to resting levels quickly after cessation of exercise (*24, 25*). Increasing brain activity results in higher energy expenditure in the axon-myelin unit (*26*) which in prolonged exercise like in a marathon might result in myelin lipid catabolism and usage of myelin-derived fatty acids for energy production.

Prolonged physical activity during a marathon results in energy shortage that may transiently impact brain structure and function. Thus, animal studies show that undernutrition impacts myelination negatively (*27, 28*). Indeed, myelin is sensitive to micro-and-macro nutrient deficiencies (*29*). Perhaps the most striking condition of dietary scarcity is anorexia nervosa, a severe psychiatric disorder, affecting predominantly women, in which body energy reserves are depleted following extreme food restriction and disturbances in cognition surrounding eating and weight (*30*). Deficits in volumes of both gray matter and white matter myelin content and connectivity are altered in anorexia nervosa as assessed by MRI studies (*31–35*). These findings open the question of whether these abnormalities may result from undernutrition and specific lipid nutritional imbalances (*32, 35*). Interestingly, the genetic architecture of anorexia nervosa shows significant genetic correlations with metabolic (including glycemic) and lipid traits (*36*), and therefore, it is conceivable that they impact in the lipid-rich myelin structure and content as well as its contribution to energy expenditure.

Physical activity is a key factor in maintaining health across the lifespan. Regular moderate-intensity training reduces oxidative stress, decreases inflammatory markers such as interleukin-6, and helps to preserve cardiovascular fitness and brain function (*37*). In contrast, strenuous, anaerobic physical activity is a risk factor for amyotrophic lateral sclerosis, as assessed by recent meta-analyses (*38, 39*). Though the risk rise (26% higher) is low in individuals with a history of vigorous physical activity, there is a five-fold mortality rate increase in professional athletes (*39*). Vigorous physical activity likely increases glutamate excitotoxicity and oxidative stress, hastening neurodegeneration in individuals with a genetic predisposition for the development of ALS (*38*). Therefore, while the impact of marathon running in well trained individuals is null, for those with ALS genetic risk, disease vulnerable motor areas are impacted more on otherwise physiological myelin expenditure (e.g., corticospinal tract), as myelin itself and myelin-producing oligodendrocytes are also vulnerable to glutamate excitotoxicity as well (*40*).

Myelin plasticity is a hallmark of brain adaptation to neuronal activity as it modifies myelin structure and increases and decreases myelination, causing thickening and thinning of the myelin sheath (*41, 42*). Our findings in marathon runners strongly suggest that widespread dramatic myelin loss, likely thinning, in exhausting conditions represents a new form of plasticity whereby brain function is preserved at the expense of myelin to optimize signal speed propagation.

### Concluding paragraph

Collectively, the findings reported here imply that the bulk of myelin is consumed upon strenuous conditions and steadily recovered later on, pointing to a novel concept of metabolic myelin plasticity. It remains to be clarified whether myelin constituents, mainly lipids, are used for fueling brain function in common life tasks including memory acquisition and retrieval, thought, sensory-motor processing, and hunger regulation, to name a few. The mechanisms driving metabolic myelin plasticity might be piloted by neural activity, and may also be modulated by neurotransmitter release at the axon-myelin unit during action potential propagation along with synaptic activity.

## Acknowledgments

We thank runner volunteers for their generosity and patience. A. Planas, J.J. Lucas, F.M. Goñi, F. Kirchhoff, G. Foffani, JP. Bolaños, A. Volterra, M.M. Panicker and F. Pérez-Cerdá for insightful discussions and suggestions on the manuscript. We also thank Osatek-Galdakao for providing human imaging facilities for MRI scans.

## Funding

Spanish Ministry of Science and Innovation and Universities grant PID2022-143020OB-I00 (CM)

Spanish Ministry of Science and Innovation and Universities grant PID2020-118546RB-I00 (PRC)

Basque Government grant IT-1551-22 (CM)

Basque Government grant KK-2021/0009 (PRC)

CIBERNED Network, Instituto Carlos III, grant CB06/05/0076 (CM)

Ikerbasque Foundation, the Ikerbasque Research Professors program (PRC)

## Author contributions

Conceptualization: PRC, CM

Methodology: PRC, AC, DP, MMG, ARA, CM

Investigation: PRC, AC, DP, MMG, ARA, CM

Funding acquisition: PRC, CM

Supervision: PRC, CM

Writing – original draft: PRC, CM

Writing –review and editing: PRC, AC, DP, MMG, ARA, CM

## Competing interests

Authors declare that they have no competing interests.

## Data and materials availability

All data, code, and materials used in this work will be available from the authors upon reasonable request (contact: Pedro Ramos-Cabrer)

## Supplementary Materials

### Materials and Methods

#### Subjects

Runners were recruited at the Donostia city marathon 2022 (n=2), and the Zegama-Aizkorri mountain marathon 2023 (n=2). All of them were well-trained men aged 45-73 years old completed the marathon in a good healthy state. Individual data are provided in table S3. All subjects provided volunteer consent prior to participation, and the experiments were conducted in accordance with the Helsinki Declaration (2001). Imaging sessions were carried out 24-48 before and after the marathon (n=4), and two weeks later (n=2). At all instances, individuals were well hydrated.

#### Data acquisition

MRI scans were acquired on a 3T whole body MRI system (Achieva Dstream, Philips Medical System, Best, The Netherlands) using the internal quadrature body coil for transmission and a 32-channel phased array coil for reception. Each subject underwent an imaging protocol that included 1) a multi echo 3D gradient and spin echo sequence (Grase) for myelin water imaging, with the following parameters: TR = 2000ms; 32 echoes with a minimum echo time of 9.3 ms and maximum of 298 ms; SENSE 2.5; flip angle 90°, bandwidth in EPI frequency direction 2591; Field of View (FOV) 230 mm^2^; 78 slices in transverse orientation; voxel size 1.2 x 1.2 x 1.8 mm; total scan time 7:08 min. And 2) A High resolution T1w anatomical image in a sagittal orientation with the following parameters: TR=7.4 ms; TE=3.4 ms; matrix size 228 x 228; flip angle 9°; FOV 250 x 250 X180 mm; slice thickness 1.1 m; 300 slices, acquisition time 4:55 m.

#### Image Processing

Raw data was exported in DICOM format for off-line processing. First, data was converted into NIFTI format using dcm2niix (*43*). Anatomic brain images (skull stripped out) were obtained from T1weighted images applying BET (*44*) implementation from FSL 6.0 distribution (*45*). Population distributions of T2 values were computed voxelwise from multi spin-echo data using the Julia implementation DECAES software (*6*). For the analysis of inter individual and intraindividual changes in MWF maps, all MWF maps for subjects at different imaging sessions (pre-, post-exercise, an after recovery) were registered to a T2 weighted anatomical image of the brain of subject 1 (echo 8 of the echo train, with TE=74.48 ms), at session 1, using ANTs Syn registration (*46*). For Image segmentation and assessment of MWF values in different regions of the brain, T1Weighted images were non-linearly registered to the JHU DTI-based white-matter atlas, for white matter tracks segmentation, (*9*) and in parallel, to the LPBA40 collection of the SRI24 brain atlas, (*13*), for segmentation of gray matter regions. Transformation matrices were later applied to the MWF maps, previously linearly registered to the corresponding T1W image. Further image processing for the creation of figures has been performed with the free available software FIJI (Image-J 2.0) and ITK-Snap 4.0.1.

### Technical considerations and limitations

#### Multiecho T2 sequences allow differentiation between water with different T2 relaxation times

Water in the brain is located in the cerebrospinal fluid (CSF), extracellular space, intracellular space and in between myelin layers. Around 70-85% of the brain mass is water, and approximately 40% of the mass in myelinated axonal tracts is compartmentalized water.

In human white matter, MRI signal from water can be separated into three pools based on its T2 relaxation time. The longest T2 component, around 2000 ms, is due to cerebrospinal fluid. The intermediate component, around 100 ms, arises both from intracellular and extracellular water; and the shortest T2 component, around 20 ms is due to water trapped between myelin bilayers (or myelin water fraction-MWF). CSF is often neglected in myelin imaging with MRI because of the low amount of CSF in WM (*47*). MWF is defined as the ratio of the area of short T2 components to the area of all T2 components (*48, 49*).

T2 mapping is a well stablished method for myelin quantification. Originally a T2 multi-echo spin-echo sequence was developed but the main drawback was the long scanning times which prevented its application in clinical scenarios. In the last years, together with other methods, a combined gradient and spin echo (GRASE) sequence has been adopted for myelin water imaging (MWI), reducing the acquisition time to less than 10 min with full brain coverage (*50*). Reproducibility of MWI with this technique has been demonstrated in a multi-site multi-vendor study (*51*) and with different reconstruction methods (*52*).

However, MWI with T2 mapping is not without challenges and several factors need to be considered. Signal to noise ratio in MWI with T2 mapping is low. Myelin signal decays rapidly as the MWF is approximately 10%, and myelin water’s relaxation time is relatively short at 10 ms, which makes changes in MWF difficult to detect (*53*).

#### MWF confounding factors

A possible confounding effect on assimilating MWF to myelin content is brain edema, which may occur in strenuous exercise like high mountain ultramarathons but has not been observed in 42.195 m marathons (*54, 55*). Moreover, edema/inflammation during new lesion formation in multiple sclerosis has a minor impact, if any, on MWF (*56*). These previous reports indicate that even if minor edema appears in a standard marathon run, the impact in MWF would be marginal.

Another confounding factor can be iron, the main source of paramagnetic susceptibility in the brain (*57*). Iron shortens T2 components and thus, potentially increases MWF values (*58*). Thus, total iron depletion in experimental animals reduces MWF values by a fourth (*58*). However, serum iron levels increase or remain similar after ultramarathon and marathon running, respectively (*59, 60*). Therefore, it is unlikely that changes in iron homeostasis contribute significantly to MWF reduction after prolonged endurance exercise, and if so, it may underestimate real myelin content reduction.

Lastly fiber orientation may also affect myelin water T2, as well as the intracellular and extracellular water T2; however, MWF has a weaker innate orientation dependence (*61*). Moreover, MWF value comparisons within and among subjects at all stages were made at similar MRI plane levels and orientation.

---

**Supplementary Figure S1.**
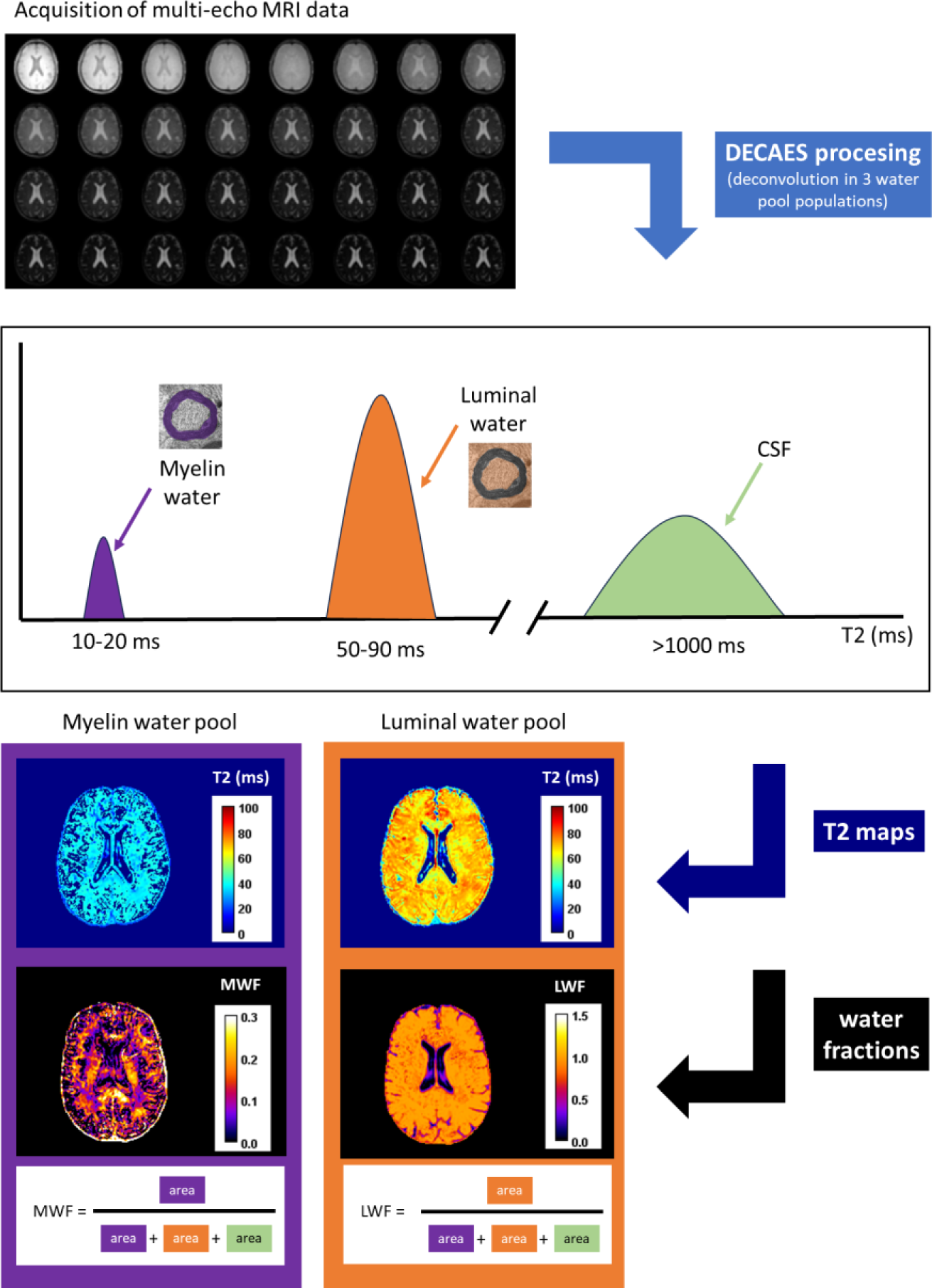
Schematic representation of the DECAES procedure to extract myelin water fraction values. First, an MRI study is conducted to collect multi-echo images that were processed using decomposition and component analysis of exponential signals (DECAES). This procedure uses the relaxation curves in each pixel to deconvolve the signal from 3 different pools of water; 1) water protons in myelin sheaths (short T2 values), 2) inter- and intra-cellular water molecules (luminal water fraction, with intermediate T2 values), and 3) signal from CSF (unrestricted water pool, with very large T2 values). Thus, a set of two T2 maps (short T2 population and intermediate T2 population, the CSF contribution is generally obviated) are generated, along with two maps of water fraction (% of contribution of that fraction to the total signal), one of them reflecting the Myelin Water Fraction (MWF) and the other one the luminal water fraction (LWF). The approach used in this study computes MWF deriving from the area of myelin protons without differentiating luminal protons and unrestricted water pools whose combined area is estimated, as established by Doucette et al. (*6*), MacKay et al. (*7*), Caverzasi et al. (*8*). ms, miliseconds.

**Supplementary Figure S2.**
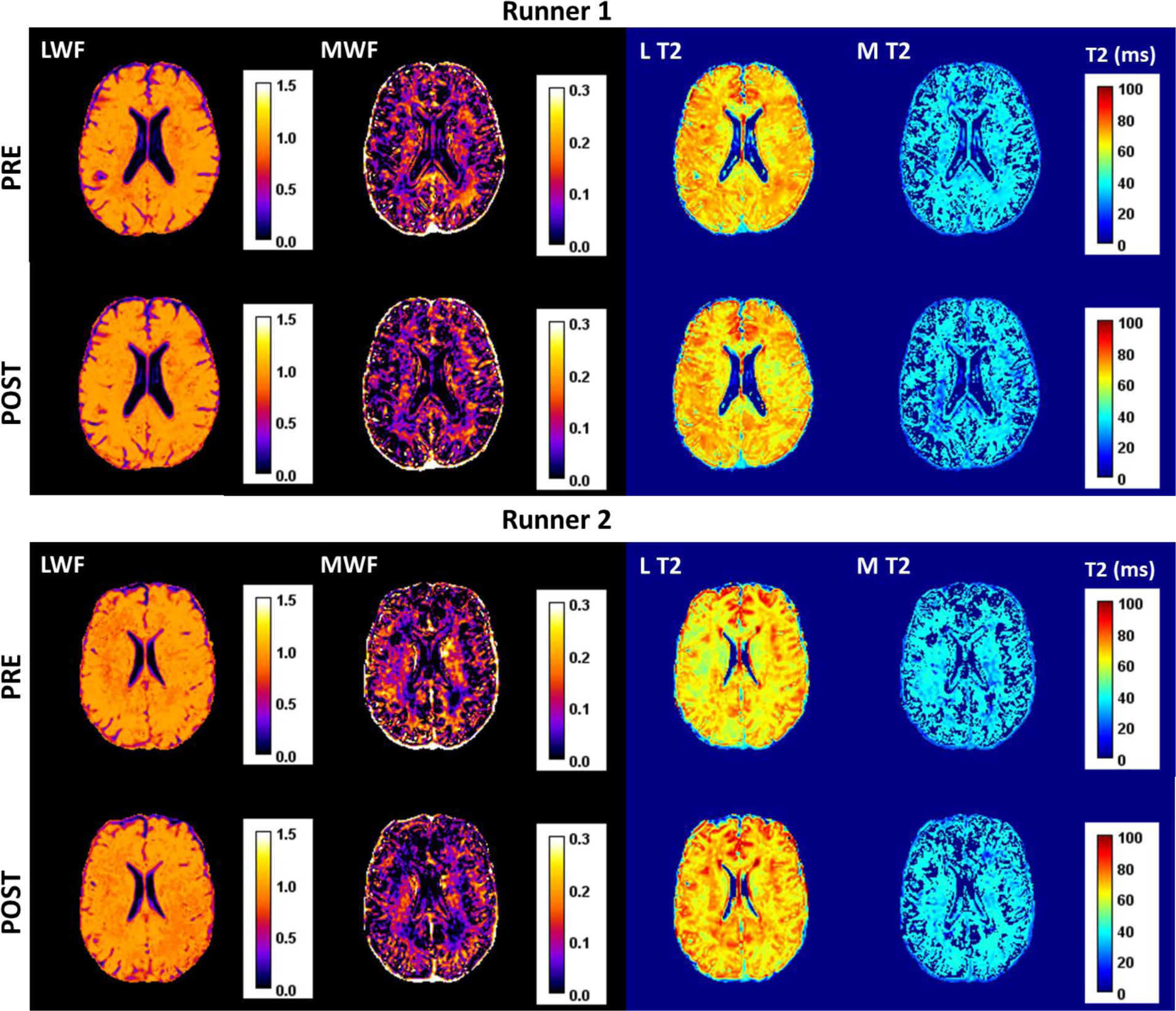
Individual results from the Multicomponent Relaxometry Magnetic Resonance Imaging Analysis of Subjects 1 and 2. From left to right representative parametric maps of: Luminal Water Fraction (LWF), Myelin Water Fraction (MWF), Luminal T2 relaxation times (L T2) and Myelin T2 relaxation times (M T2), are presented for the different subjects at the different imaging sessions. Only WMF shows visible changes over sessions.

**Supplementary Figure S3.**
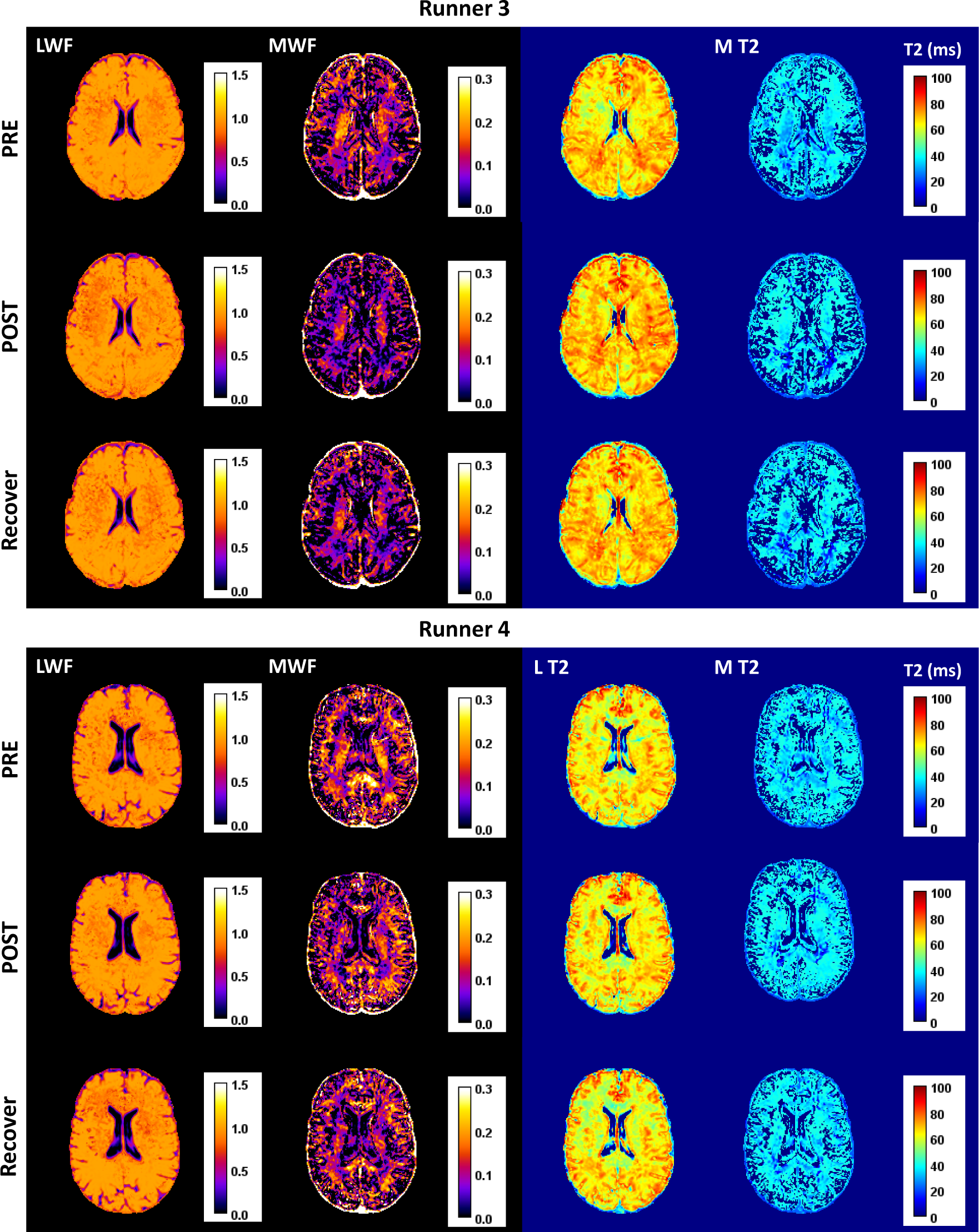
Individual results from the Multicomponent Relaxometry Magnetic Resonance Imaging Analysis of Subjects 3 and 4. From left to right representative parametric maps of: Luminal Water Fraction (LWF), Myelin Water Fraction (MWF), Luminal T2 relaxation times (L T2) and Myelin T2 relaxation times (M T2), are presented for the different subjects at the different imaging sessions. Only WMF shows visible changes over sessions.

**Figure S4.**
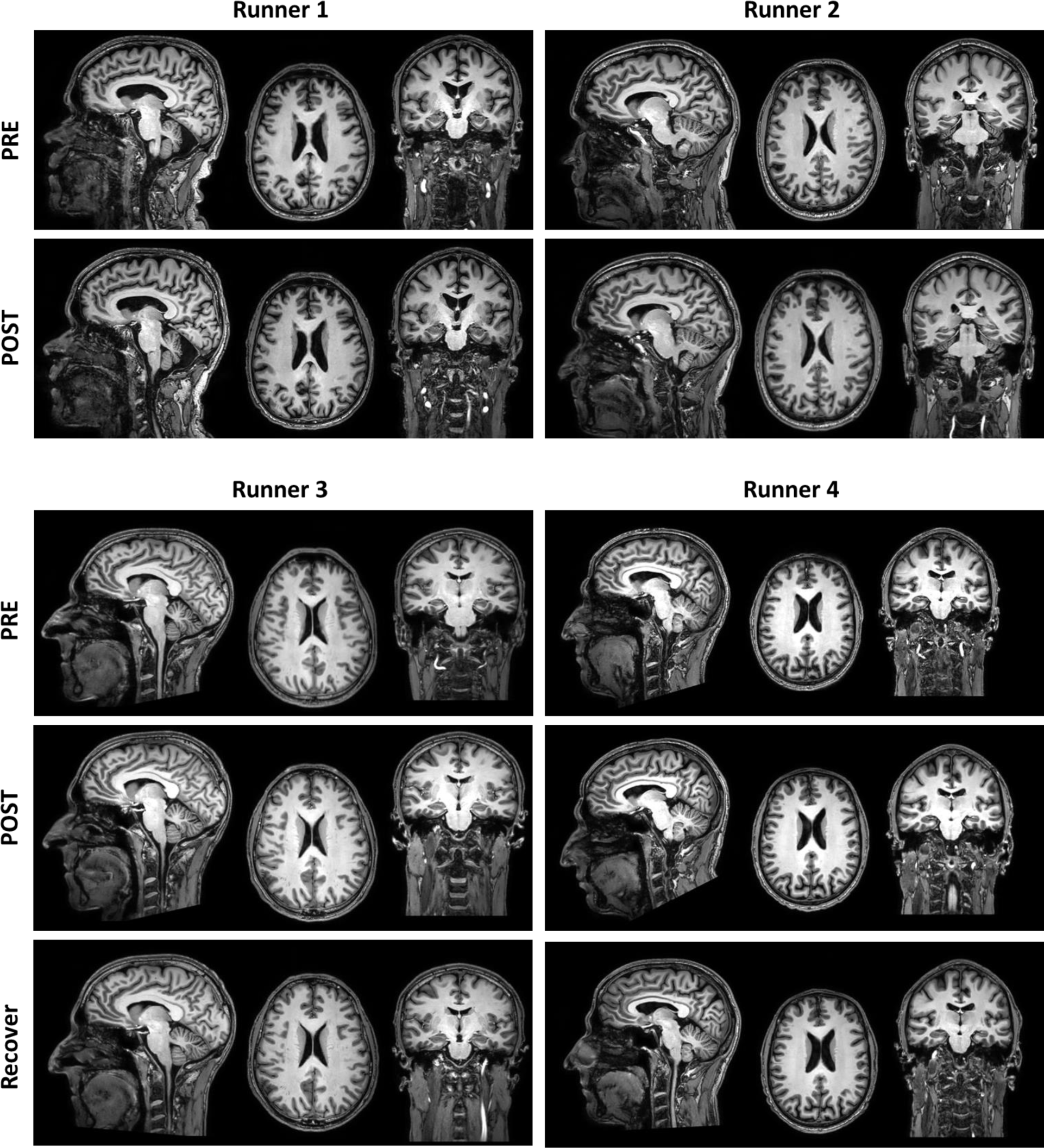
Hight resolution 3D T1 weighted MRI images of all subjects. Sagittal, axial and coronal views of the 3D images acquired with T1 weighting for anatomical reference, pre- and post-exercise and post recovery for subjects 3 and 4. Images show no significant interindividual differences for the different imaging sessions.

**Table S1.** Volumes of brain regions across subjects and sessions. Total brain volumes include CSF, gray matter, white matter and deep brain, brainstem and cerebellum.

| Table S1. Volumes of brain regions across subjects and sessions. |  |  |  |  |  |  |  |
| --- | --- | --- | --- | --- | --- | --- | --- |
| Total brain volumes include CSF, gray matter, white matter and deep brain, brainstem and cerebellum. |  |  |  |  |  |  |  |
| Volume (cm<sup>3</sup>) |  |  |  |  |  |  |  |
|  | Total brain | CSF | Gray matter | White matter | Deep brain | Brainstem | Cerebellum |
| **Runner 1** |  |  |  |  |  |  |  |
| pre-run | 1430 | 290 | 610 | 483 | 47 | 23 | 138 |
| post-run | 1432 | 291 | 609 | 485 | 46 | 23 | 139 |
| **Runner 2** |  |  |  |  |  |  |  |
| pre-run | 1269 | 223 | 559 | 440 | 47 | 24 | 155 |
| post-run | 1273 | 220 | 561 | 445 | 47 | 24 | 154 |
| **Runner 3** |  |  |  |  |  |  |  |
| pre-run | 1420 | 235 | 621 | 516 | 49 | 23 | 147 |
| post-run | 1428 | 232 | 623 | 524 | 48 | 24 | 148 |
| 2-weeks recovery | 1417 | 231 | 616 | 522 | 48 | 24 | 146 |
| **Runner 4** |  |  |  |  |  |  |  |
| pre-run | 1260 | 235 | 541 | 444 | 40 | 19 | 131 |
| post-run | 1280 | 239 | 563 | 435 | 42 | 19 | 133 |
| 2-weeks recovery | 1276 | 238 | 562 | 435 | 42 | 19 | 133 |

**Table S2.** Myelin Water Fraction (MWF) row data. Raw MWF values for all subjects in all imaging sessions for each region of the brain using the two atlases (JHU and SRI24), presented as mean and standard deviation in the specific region

| GRAY MATTER | Subject 1 |  |  |  | Subject 2 |  |  |  | Subject 3 |  |  |  |  |  | Subject 4 |  |  |  |  |  |
| --- | --- | --- | --- | --- | --- | --- | --- | --- | --- | --- | --- | --- | --- | --- | --- | --- | --- | --- | --- | --- |
|  | Session 1 |  | Session 2 |  | Session 1 |  | Session 2 |  | Session 1 |  | Session 2 |  | Session 3 |  | Session 1 |  | Session 2 |  | Session 3 |  |
| Region | Mean | STD | Mean | STD | Mean | STD | Mean | STD | Mean | STD | Mean | STD | Mean | STD | Mean | STD | Mean | STD | Mean | STD |
| background | 0.012 | 0.049 | 0.011 | 0.047 | 0.011 | 0.047 | 0.009 | 0.041 | 0.010 | 0.043 | 0.009 | 0.040 | 0.010 | 0.042 | 0.011 | 0.047 | 0.011 | 0.045 | 0.011 | 0.046 |
| L_superior_frontal_gyrus | 0.076 | 0.054 | 0.061 | 0.054 | 0.075 | 0.065 | 0.050 | 0.055 | 0.067 | 0.056 | 0.057 | 0.061 | 0.064 | 0.060 | 0.052 | 0.061 | 0.058 | 0.055 | 0.062 | 0.061 |
| R_superior_frontal_gyrus | 0.071 | 0.058 | 0.066 | 0.056 | 0.068 | 0.064 | 0.054 | 0.056 | 0.062 | 0.061 | 0.049 | 0.055 | 0.059 | 0.062 | 0.064 | 0.066 | 0.054 | 0.055 | 0.064 | 0.060 |
| L_middle_frontal_gyrus | 0.070 | 0.049 | 0.051 | 0.043 | 0.081 | 0.049 | 0.041 | 0.043 | 0.087 | 0.051 | 0.047 | 0.045 | 0.072 | 0.048 | 0.052 | 0.057 | 0.048 | 0.048 | 0.062 | 0.054 |
| R_middle_frontal_gyrus | 0.055 | 0.050 | 0.064 | 0.050 | 0.064 | 0.052 | 0.060 | 0.046 | 0.059 | 0.054 | 0.043 | 0.046 | 0.050 | 0.047 | 0.075 | 0.061 | 0.071 | 0.051 | 0.076 | 0.056 |
| L_inferior_frontal_gyrus | 0.058 | 0.053 | 0.039 | 0.044 | 0.060 | 0.053 | 0.049 | 0.060 | 0.076 | 0.067 | 0.035 | 0.052 | 0.057 | 0.059 | 0.060 | 0.065 | 0.056 | 0.058 | 0.059 | 0.057 |
| R_inferior_frontal_gyrus | 0.062 | 0.048 | 0.077 | 0.046 | 0.060 | 0.063 | 0.064 | 0.060 | 0.055 | 0.059 | 0.050 | 0.058 | 0.045 | 0.051 | 0.108 | 0.059 | 0.096 | 0.050 | 0.096 | 0.048 |
| L_precentral_gyrus | 0.110 | 0.059 | 0.086 | 0.060 | 0.128 | 0.068 | 0.079 | 0.065 | 0.103 | 0.055 | 0.092 | 0.063 | 0.088 | 0.057 | 0.089 | 0.076 | 0.098 | 0.066 | 0.098 | 0.070 |
| R_precentral_gyrus | 0.091 | 0.055 | 0.074 | 0.052 | 0.117 | 0.064 | 0.087 | 0.051 | 0.084 | 0.062 | 0.079 | 0.062 | 0.076 | 0.063 | 0.098 | 0.068 | 0.091 | 0.051 | 0.099 | 0.058 |
| L_middle_orbitofrontal_gyrus | 0.105 | 0.087 | 0.093 | 0.086 | 0.057 | 0.061 | 0.073 | 0.090 | 0.074 | 0.084 | 0.042 | 0.073 | 0.048 | 0.073 | 0.084 | 0.077 | 0.057 | 0.063 | 0.070 | 0.065 |
| R_middle_orbitofrontal_gyrus | 0.103 | 0.089 | 0.111 | 0.088 | 0.068 | 0.080 | 0.077 | 0.087 | 0.098 | 0.094 | 0.069 | 0.094 | 0.069 | 0.086 | 0.114 | 0.074 | 0.103 | 0.072 | 0.110 | 0.065 |
| L_lateral_orbitofrontal_gyrus | 0.090 | 0.076 | 0.087 | 0.068 | 0.056 | 0.071 | 0.069 | 0.086 | 0.070 | 0.099 | 0.038 | 0.078 | 0.052 | 0.092 | 0.085 | 0.074 | 0.065 | 0.075 | 0.071 | 0.070 |
| R_lateral_orbitofrontal_gyrus | 0.085 | 0.063 | 0.094 | 0.069 | 0.076 | 0.094 | 0.089 | 0.088 | 0.075 | 0.093 | 0.061 | 0.092 | 0.049 | 0.077 | 0.137 | 0.059 | 0.121 | 0.062 | 0.111 | 0.063 |
| L_gyrus_rectus | 0.094 | 0.062 | 0.091 | 0.071 | 0.074 | 0.092 | 0.055 | 0.081 | 0.048 | 0.073 | 0.036 | 0.064 | 0.031 | 0.048 | 0.110 | 0.069 | 0.079 | 0.070 | 0.065 | 0.072 |
| R_gyrus_rectus | 0.074 | 0.062 | 0.081 | 0.059 | 0.046 | 0.050 | 0.051 | 0.066 | 0.042 | 0.076 | 0.028 | 0.067 | 0.035 | 0.065 | 0.118 | 0.068 | 0.078 | 0.066 | 0.054 | 0.063 |
| L_postcentral_gyrus | 0.100 | 0.074 | 0.075 | 0.077 | 0.120 | 0.057 | 0.086 | 0.059 | 0.097 | 0.059 | 0.087 | 0.060 | 0.083 | 0.059 | 0.096 | 0.068 | 0.092 | 0.061 | 0.090 | 0.064 |
| R_postcentral_gyrus | 0.073 | 0.053 | 0.060 | 0.052 | 0.107 | 0.056 | 0.090 | 0.047 | 0.090 | 0.069 | 0.081 | 0.062 | 0.074 | 0.060 | 0.115 | 0.065 | 0.095 | 0.057 | 0.086 | 0.061 |
| L_superior_parietal_gyrus | 0.067 | 0.050 | 0.037 | 0.050 | 0.084 | 0.051 | 0.057 | 0.049 | 0.078 | 0.081 | 0.067 | 0.082 | 0.059 | 0.082 | 0.051 | 0.049 | 0.083 | 0.059 | 0.057 | 0.059 |
| R_superior_parietal_gyrus | 0.063 | 0.061 | 0.048 | 0.068 | 0.072 | 0.051 | 0.059 | 0.049 | 0.080 | 0.091 | 0.063 | 0.082 | 0.058 | 0.086 | 0.080 | 0.060 | 0.073 | 0.055 | 0.051 | 0.054 |
| L_supramarginal_gyrus | 0.062 | 0.060 | 0.034 | 0.046 | 0.054 | 0.057 | 0.050 | 0.064 | 0.060 | 0.073 | 0.048 | 0.068 | 0.042 | 0.059 | 0.049 | 0.063 | 0.050 | 0.065 | 0.045 | 0.061 |
| R_supramarginal_gyrus | 0.042 | 0.046 | 0.036 | 0.048 | 0.063 | 0.061 | 0.061 | 0.054 | 0.041 | 0.059 | 0.045 | 0.069 | 0.028 | 0.049 | 0.074 | 0.058 | 0.052 | 0.051 | 0.046 | 0.049 |
| L angular_gyrus | 0.068 | 0.087 | 0.026 | 0.048 | 0.044 | 0.050 | 0.061 | 0.059 | 0.039 | 0.063 | 0.045 | 0.068 | 0.038 | 0.067 | 0.031 | 0.044 | 0.054 | 0.063 | 0.050 | 0.066 |
| R angular_gyrus | 0.048 | 0.051 | 0.035 | 0.048 | 0.052 | 0.051 | 0.084 | 0.055 | 0.034 | 0.065 | 0.041 | 0.070 | 0.024 | 0.051 | 0.064 | 0.048 | 0.042 | 0.049 | 0.030 | 0.042 |
| L_precuneus | 0.093 | 0.080 | 0.067 | 0.081 | 0.070 | 0.066 | 0.055 | 0.058 | 0.077 | 0.116 | 0.055 | 0.098 | 0.056 | 0.102 | 0.063 | 0.062 | 0.087 | 0.060 | 0.039 | 0.052 |
| R_precuneus | 0.119 | 0.119 | 0.089 | 0.096 | 0.065 | 0.070 | 0.064 | 0.080 | 0.102 | 0.123 | 0.057 | 0.095 | 0.078 | 0.115 | 0.068 | 0.070 | 0.084 | 0.063 | 0.051 | 0.067 |
| L_superior_occipital_gyrus | 0.134 | 0.112 | 0.095 | 0.121 | 0.099 | 0.062 | 0.095 | 0.075 | 0.092 | 0.120 | 0.093 | 0.126 | 0.087 | 0.126 | 0.071 | 0.050 | 0.101 | 0.071 | 0.078 | 0.079 |
| R_superior_occipital_gyrus | 0.135 | 0.113 | 0.091 | 0.106 | 0.102 | 0.072 | 0.113 | 0.067 | 0.091 | 0.121 | 0.079 | 0.110 | 0.075 | 0.108 | 0.092 | 0.052 | 0.089 | 0.078 | 0.094 | 0.090 |
| L_middle_occipital_gyrus | 0.104 | 0.089 | 0.052 | 0.076 | 0.076 | 0.070 | 0.093 | 0.068 | 0.053 | 0.073 | 0.054 | 0.067 | 0.044 | 0.066 | 0.064 | 0.051 | 0.095 | 0.064 | 0.061 | 0.070 |
| R_middle_occipital_gyrus | 0.109 | 0.082 | 0.097 | 0.081 | 0.104 | 0.076 | 0.130 | 0.068 | 0.069 | 0.099 | 0.080 | 0.096 | 0.068 | 0.092 | 0.117 | 0.056 | 0.105 | 0.065 | 0.092 | 0.071 |
| L_inferior_occipital_gyrus | 0.134 | 0.091 | 0.097 | 0.102 | 0.134 | 0.102 | 0.135 | 0.080 | 0.104 | 0.102 | 0.075 | 0.078 | 0.070 | 0.086 | 0.097 | 0.057 | 0.122 | 0.064 | 0.102 | 0.101 |
| R_inferior_occipital_gyrus | 0.186 | 0.134 | 0.135 | 0.144 | 0.209 | 0.135 | 0.187 | 0.134 | 0.158 | 0.141 | 0.138 | 0.115 | 0.126 | 0.113 | 0.143 | 0.067 | 0.167 | 0.090 | 0.180 | 0.122 |
| L_cuneus | 0.212 | 0.179 | 0.190 | 0.195 | 0.150 | 0.068 | 0.109 | 0.066 | 0.082 | 0.114 | 0.102 | 0.122 | 0.082 | 0.109 | 0.121 | 0.058 | 0.153 | 0.087 | 0.123 | 0.102 |
| R_cuneus | 0.225 | 0.199 | 0.145 | 0.162 | 0.151 | 0.124 | 0.168 | 0.151 | 0.169 | 0.188 | 0.135 | 0.135 | 0.151 | 0.144 | 0.123 | 0.071 | 0.139 | 0.097 | 0.139 | 0.114 |
| L_superior_temporal_gyrus | 0.065 | 0.069 | 0.051 | 0.059 | 0.042 | 0.062 | 0.053 | 0.069 | 0.048 | 0.063 | 0.032 | 0.056 | 0.039 | 0.061 | 0.062 | 0.078 | 0.047 | 0.062 | 0.059 | 0.069 |
| R_superior_temporal_gyrus | 0.075 | 0.069 | 0.082 | 0.066 | 0.064 | 0.074 | 0.053 | 0.062 | 0.048 | 0.066 | 0.044 | 0.069 | 0.042 | 0.069 | 0.118 | 0.082 | 0.087 | 0.062 | 0.088 | 0.063 |
| L_middle_temporal_gyrus | 0.051 | 0.066 | 0.043 | 0.054 | 0.040 | 0.061 | 0.061 | 0.059 | 0.034 | 0.055 | 0.022 | 0.047 | 0.021 | 0.042 | 0.045 | 0.061 | 0.039 | 0.051 | 0.046 | 0.058 |
| R_middle_temporal_gyrus | 0.068 | 0.056 | 0.058 | 0.053 | 0.080 | 0.069 | 0.060 | 0.057 | 0.047 | 0.061 | 0.037 | 0.059 | 0.032 | 0.049 | 0.122 | 0.073 | 0.088 | 0.064 | 0.082 | 0.063 |
| L_inferior_temporal_gyrus | 0.069 | 0.109 | 0.060 | 0.091 | 0.040 | 0.071 | 0.049 | 0.062 | 0.040 | 0.069 | 0.027 | 0.064 | 0.034 | 0.069 | 0.049 | 0.064 | 0.053 | 0.071 | 0.048 | 0.070 |
| R_inferior_temporal_gyrus | 0.101 | 0.121 | 0.090 | 0.108 | 0.096 | 0.104 | 0.075 | 0.108 | 0.052 | 0.089 | 0.038 | 0.070 | 0.039 | 0.068 | 0.093 | 0.077 | 0.079 | 0.078 | 0.088 | 0.087 |
| L_parahippocampal_gyrus | 0.093 | 0.109 | 0.088 | 0.091 | 0.099 | 0.119 | 0.074 | 0.111 | 0.071 | 0.104 | 0.056 | 0.094 | 0.066 | 0.103 | 0.092 | 0.109 | 0.093 | 0.098 | 0.079 | 0.101 |
| R_parahippocampal_gyrus | 0.118 | 0.111 | 0.097 | 0.101 | 0.124 | 0.123 | 0.109 | 0.134 | 0.091 | 0.108 | 0.063 | 0.100 | 0.075 | 0.106 | 0.130 | 0.097 | 0.102 | 0.093 | 0.132 | 0.107 |
| L lingual_gyrus | 0.158 | 0.098 | 0.127 | 0.122 | 0.165 | 0.101 | 0.122 | 0.107 | 0.086 | 0.097 | 0.091 | 0.095 | 0.077 | 0.094 | 0.141 | 0.066 | 0.142 | 0.062 | 0.108 | 0.078 |
| R lingual_gyrus | 0.155 | 0.113 | 0.120 | 0.104 | 0.157 | 0.088 | 0.116 | 0.120 | 0.104 | 0.122 | 0.090 | 0.099 | 0.094 | 0.103 | 0.156 | 0.062 | 0.147 | 0.081 | 0.135 | 0.087 |
| L_fusiform_gyrus | 0.077 | 0.081 | 0.057 | 0.072 | 0.065 | 0.071 | 0.053 | 0.069 | 0.052 | 0.077 | 0.033 | 0.058 | 0.036 | 0.063 | 0.056 | 0.070 | 0.064 | 0.065 | 0.045 | 0.075 |
| R_fusiform_gyrus | 0.084 | 0.097 | 0.080 | 0.075 | 0.117 | 0.093 | 0.058 | 0.078 | 0.059 | 0.079 | 0.034 | 0.057 | 0.041 | 0.064 | 0.106 | 0.075 | 0.094 | 0.076 | 0.086 | 0.088 |
| L_insular_cortex | 0.044 | 0.050 | 0.039 | 0.048 | 0.038 | 0.065 | 0.021 | 0.042 | 0.026 | 0.049 | 0.013 | 0.044 | 0.012 | 0.036 | 0.028 | 0.048 | 0.017 | 0.043 | 0.019 | 0.041 |
| R_insular_cortex | 0.055 | 0.068 | 0.068 | 0.059 | 0.041 | 0.071 | 0.030 | 0.064 | 0.038 | 0.072 | 0.020 | 0.054 | 0.022 | 0.062 | 0.053 | 0.068 | 0.043 | 0.063 | 0.040 | 0.058 |
| L_cingulate_gyrus | 0.065 | 0.059 | 0.049 | 0.051 | 0.049 | 0.069 | 0.036 | 0.060 | 0.060 | 0.056 | 0.028 | 0.050 | 0.030 | 0.045 | 0.054 | 0.065 | 0.056 | 0.056 | 0.043 | 0.058 |
| R_cingulate_gyrus | 0.064 | 0.074 | 0.043 | 0.055 | 0.049 | 0.068 | 0.036 | 0.059 | 0.059 | 0.060 | 0.019 | 0.038 | 0.033 | 0.051 | 0.059 | 0.067 | 0.056 | 0.056 | 0.046 | 0.063 |
| L_caudate | 0.089 | 0.051 | 0.079 | 0.050 | 0.115 | 0.086 | 0.094 | 0.070 | 0.118 | 0.057 | 0.043 | 0.052 | 0.068 | 0.059 | 0.070 | 0.057 | 0.066 | 0.050 | 0.050 | 0.051 |
| R_caudate | 0.075 | 0.045 | 0.082 | 0.050 | 0.132 | 0.089 | 0.089 | 0.083 | 0.078 | 0.062 | 0.040 | 0.053 | 0.050 | 0.055 | 0.070 | 0.064 | 0.061 | 0.061 | 0.074 | 0.054 |
| L_putamen | 0.167 | 0.094 | 0.146 | 0.091 | 0.263 | 0.125 | 0.192 | 0.112 | 0.190 | 0.111 | 0.138 | 0.107 | 0.151 | 0.116 | 0.181 | 0.112 | 0.142 | 0.104 | 0.153 | 0.114 |
| R_putamen | 0.181 | 0.076 | 0.159 | 0.077 | 0.250 | 0.129 | 0.194 | 0.129 | 0.189 | 0.102 | 0.147 | 0.105 | 0.152 | 0.109 | 0.199 | 0.137 | 0.175 | 0.113 | 0.171 | 0.110 |
| L_hippocampus | 0.049 | 0.081 | 0.042 | 0.052 | 0.036 | 0.067 | 0.027 | 0.058 | 0.020 | 0.047 | 0.009 | 0.041 | 0.014 | 0.045 | 0.048 | 0.074 | 0.042 | 0.065 | 0.037 | 0.058 |
| R_hippocampus | 0.055 | 0.070 | 0.045 | 0.059 | 0.041 | 0.054 | 0.021 | 0.053 | 0.034 | 0.054 | 0.012 | 0.036 | 0.017 | 0.044 | 0.073 | 0.062 | 0.050 | 0.055 | 0.059 | 0.062 |
| Cerebellum | 0.069 | 0.085 | 0.056 | 0.075 | 0.060 | 0.061 | 0.046 | 0.061 | 0.079 | 0.096 | 0.049 | 0.082 | 0.053 | 0.078 | 0.084 | 0.075 | 0.085 | 0.075 | 0.068 | 0.075 |
| Brainstem | 0.131 | 0.066 | 0.119 | 0.067 | 0.125 | 0.087 | 0.096 | 0.080 | 0.123 | 0.069 | 0.088 | 0.069 | 0.114 | 0.073 | 0.149 | 0.085 | 0.130 | 0.069 | 0.128 | 0.074 |

| WHITE MATTER | Subject 1 |  |  |  | Subject 2 |  |  |  | Subject 3 |  |  |  |  |  | Subject 4 |  |  |  |  |  |
| --- | --- | --- | --- | --- | --- | --- | --- | --- | --- | --- | --- | --- | --- | --- | --- | --- | --- | --- | --- | --- |
|  | Session 1 |  | Session 2 |  | Session 1 |  | Session 2 |  | Session 1 |  | Session 2 |  | Session 3 |  | Session 1 |  | Session 2 |  | Session 3 |  |
| Region | Mean | STD | Mean | STD | Mean | STD | Mean | STD | Mean | STD | Mean | STD | Mean | STD | Mean | STD | Mean | STD | Mean | STD |
| background | 0.022 | 0.061 | 0.019 | 0.056 | 0.020 | 0.057 | 0.017 | 0.051 | 0.019 | 0.056 | 0.015 | 0.050 | 0.016 | 0.051 | 0.020 | 0.055 | 0.019 | 0.054 | 0.019 | 0.054 |
| Middle cerebellar peduncle | 0.104 | 0.050 | 0.091 | 0.053 | 0.089 | 0.044 | 0.062 | 0.037 | 0.110 | 0.046 | 0.080 | 0.045 | 0.098 | 0.047 | 0.134 | 0.050 | 0.103 | 0.053 | 0.113 | 0.046 |
| Pontine crossing tract | 0.148 | 0.039 | 0.153 | 0.035 | 0.121 | 0.027 | 0.102 | 0.028 | 0.125 | 0.033 | 0.105 | 0.026 | 0.106 | 0.032 | 0.142 | 0.031 | 0.138 | 0.036 | 0.114 | 0.036 |
| Genu of corpus callosum | 0.102 | 0.050 | 0.095 | 0.052 | 0.095 | 0.056 | 0.067 | 0.052 | 0.096 | 0.049 | 0.050 | 0.042 | 0.062 | 0.044 | 0.127 | 0.070 | 0.107 | 0.071 | 0.132 | 0.065 |
| Body of corpus callosum | 0.088 | 0.039 | 0.067 | 0.037 | 0.069 | 0.043 | 0.050 | 0.036 | 0.078 | 0.040 | 0.040 | 0.034 | 0.057 | 0.034 | 0.083 | 0.041 | 0.072 | 0.041 | 0.056 | 0.048 |
| Splenium of corpus callosum | 0.126 | 0.059 | 0.090 | 0.051 | 0.104 | 0.061 | 0.089 | 0.060 | 0.095 | 0.041 | 0.051 | 0.039 | 0.086 | 0.042 | 0.168 | 0.068 | 0.114 | 0.055 | 0.131 | 0.064 |
| Fornix (column and body) | 0.079 | 0.042 | 0.068 | 0.035 | 0.045 | 0.046 | 0.052 | 0.037 | 0.037 | 0.042 | 0.014 | 0.027 | 0.012 | 0.025 | 0.070 | 0.050 | 0.056 | 0.043 | 0.034 | 0.038 |
| R Corticospinal tract | 0.118 | 0.037 | 0.122 | 0.046 | 0.119 | 0.034 | 0.097 | 0.024 | 0.138 | 0.037 | 0.114 | 0.031 | 0.135 | 0.036 | 0.158 | 0.038 | 0.112 | 0.043 | 0.144 | 0.035 |
| L Corticospinal tract | 0.142 | 0.042 | 0.139 | 0.044 | 0.132 | 0.039 | 0.120 | 0.038 | 0.146 | 0.025 | 0.111 | 0.032 | 0.127 | 0.028 | 0.146 | 0.033 | 0.097 | 0.034 | 0.125 | 0.032 |
| R Medial lemniscus | 0.108 | 0.036 | 0.111 | 0.037 | 0.089 | 0.043 | 0.076 | 0.027 | 0.100 | 0.045 | 0.071 | 0.029 | 0.100 | 0.036 | 0.122 | 0.037 | 0.119 | 0.049 | 0.108 | 0.032 |
| L Medial lemniscus | 0.141 | 0.053 | 0.136 | 0.048 | 0.081 | 0.038 | 0.082 | 0.036 | 0.118 | 0.044 | 0.076 | 0.036 | 0.114 | 0.039 | 0.132 | 0.042 | 0.124 | 0.042 | 0.107 | 0.041 |
| R Inferior cerebellar peduncle | 0.110 | 0.047 | 0.089 | 0.046 | 0.098 | 0.031 | 0.074 | 0.038 | 0.091 | 0.042 | 0.066 | 0.039 | 0.100 | 0.042 | 0.155 | 0.036 | 0.111 | 0.033 | 0.119 | 0.034 |
| L Inferior cerebellar peduncle | 0.144 | 0.078 | 0.100 | 0.062 | 0.078 | 0.038 | 0.061 | 0.044 | 0.102 | 0.037 | 0.074 | 0.031 | 0.102 | 0.037 | 0.133 | 0.055 | 0.101 | 0.053 | 0.103 | 0.049 |
| R Superior cerebellar peduncle | 0.138 | 0.042 | 0.121 | 0.040 | 0.137 | 0.045 | 0.096 | 0.038 | 0.125 | 0.039 | 0.096 | 0.039 | 0.121 | 0.041 | 0.178 | 0.028 | 0.154 | 0.033 | 0.161 | 0.036 |
| L Superior cerebellar peduncle | 0.133 | 0.044 | 0.115 | 0.039 | 0.112 | 0.046 | 0.110 | 0.047 | 0.141 | 0.046 | 0.091 | 0.036 | 0.119 | 0.041 | 0.150 | 0.043 | 0.140 | 0.037 | 0.121 | 0.036 |
| R Cerebral peduncle | 0.189 | 0.071 | 0.174 | 0.085 | 0.284 | 0.125 | 0.254 | 0.111 | 0.232 | 0.071 | 0.185 | 0.065 | 0.227 | 0.073 | 0.287 | 0.080 | 0.259 | 0.085 | 0.261 | 0.076 |
| L Cerebral peduncle | 0.192 | 0.062 | 0.179 | 0.079 | 0.281 | 0.129 | 0.223 | 0.104 | 0.222 | 0.067 | 0.184 | 0.069 | 0.236 | 0.073 | 0.282 | 0.095 | 0.238 | 0.084 | 0.261 | 0.086 |
| R Anterior limb of int. capsule | 0.117 | 0.066 | 0.106 | 0.067 | 0.215 | 0.106 | 0.161 | 0.102 | 0.141 | 0.067 | 0.097 | 0.069 | 0.101 | 0.070 | 0.177 | 0.108 | 0.140 | 0.079 | 0.141 | 0.085 |
| L Anterior limb of int. capsule | 0.118 | 0.070 | 0.108 | 0.053 | 0.210 | 0.121 | 0.150 | 0.103 | 0.152 | 0.065 | 0.109 | 0.066 | 0.122 | 0.076 | 0.169 | 0.091 | 0.131 | 0.077 | 0.132 | 0.086 |
| R Posterior limb of int.capsule | 0.157 | 0.041 | 0.127 | 0.049 | 0.208 | 0.082 | 0.167 | 0.079 | 0.167 | 0.044 | 0.146 | 0.053 | 0.158 | 0.053 | 0.245 | 0.061 | 0.179 | 0.058 | 0.179 | 0.066 |
| L Posterior limb of int capsule | 0.175 | 0.046 | 0.134 | 0.051 | 0.209 | 0.098 | 0.182 | 0.081 | 0.208 | 0.050 | 0.159 | 0.059 | 0.185 | 0.051 | 0.230 | 0.076 | 0.176 | 0.069 | 0.192 | 0.084 |
| R Retrolenticular part of internal capsule | 0.100 | 0.052 | 0.091 | 0.045 | 0.109 | 0.055 | 0.059 | 0.040 | 0.122 | 0.039 | 0.075 | 0.043 | 0.087 | 0.047 | 0.192 | 0.032 | 0.104 | 0.034 | 0.112 | 0.042 |
| L Retrolenticular part of internal capsule | 0.134 | 0.032 | 0.102 | 0.039 | 0.109 | 0.050 | 0.098 | 0.044 | 0.156 | 0.040 | 0.093 | 0.037 | 0.136 | 0.040 | 0.173 | 0.040 | 0.116 | 0.045 | 0.126 | 0.036 |
| R Anterior corona radiata | 0.102 | 0.028 | 0.107 | 0.032 | 0.091 | 0.031 | 0.093 | 0.028 | 0.098 | 0.033 | 0.052 | 0.033 | 0.053 | 0.039 | 0.109 | 0.041 | 0.099 | 0.041 | 0.101 | 0.036 |
| L Anterior corona radiata | 0.111 | 0.028 | 0.079 | 0.033 | 0.063 | 0.030 | 0.038 | 0.029 | 0.071 | 0.036 | 0.048 | 0.027 | 0.054 | 0.036 | 0.078 | 0.041 | 0.053 | 0.033 | 0.064 | 0.037 |
| R Superior corona radiata | 0.124 | 0.027 | 0.100 | 0.031 | 0.129 | 0.030 | 0.100 | 0.024 | 0.117 | 0.028 | 0.096 | 0.029 | 0.094 | 0.029 | 0.145 | 0.042 | 0.122 | 0.040 | 0.109 | 0.035 |
| L Superior corona radiata | 0.120 | 0.025 | 0.124 | 0.026 | 0.073 | 0.032 | 0.087 | 0.031 | 0.115 | 0.046 | 0.068 | 0.032 | 0.095 | 0.037 | 0.137 | 0.036 | 0.092 | 0.045 | 0.099 | 0.038 |
| R Posterior corona radiata | 0.094 | 0.036 | 0.064 | 0.036 | 0.064 | 0.047 | 0.049 | 0.036 | 0.082 | 0.031 | 0.060 | 0.046 | 0.061 | 0.039 | 0.107 | 0.042 | 0.093 | 0.037 | 0.081 | 0.036 |
| L Posterior corona radiata | 0.099 | 0.026 | 0.097 | 0.033 | 0.076 | 0.031 | 0.074 | 0.035 | 0.115 | 0.033 | 0.067 | 0.032 | 0.087 | 0.038 | 0.110 | 0.044 | 0.092 | 0.041 | 0.101 | 0.037 |
| R Posterior thalamic radiation | 0.065 | 0.038 | 0.091 | 0.040 | 0.079 | 0.043 | 0.032 | 0.027 | 0.063 | 0.028 | 0.030 | 0.028 | 0.046 | 0.027 | 0.116 | 0.034 | 0.101 | 0.034 | 0.102 | 0.051 |
| L Posterior thalamic radiation | 0.096 | 0.027 | 0.065 | 0.035 | 0.088 | 0.041 | 0.079 | 0.035 | 0.091 | 0.024 | 0.049 | 0.024 | 0.081 | 0.023 | 0.111 | 0.032 | 0.086 | 0.036 | 0.093 | 0.033 |
| R Sagittal stratum | 0.070 | 0.027 | 0.087 | 0.038 | 0.090 | 0.033 | 0.034 | 0.028 | 0.053 | 0.031 | 0.020 | 0.025 | 0.023 | 0.024 | 0.158 | 0.037 | 0.098 | 0.030 | 0.091 | 0.040 |
| L Sagittal stratum | 0.103 | 0.037 | 0.082 | 0.039 | 0.069 | 0.037 | 0.066 | 0.029 | 0.077 | 0.032 | 0.034 | 0.028 | 0.069 | 0.027 | 0.126 | 0.035 | 0.068 | 0.039 | 0.109 | 0.039 |
| R External capsule | 0.078 | 0.056 | 0.077 | 0.058 | 0.135 | 0.077 | 0.068 | 0.059 | 0.099 | 0.052 | 0.071 | 0.055 | 0.056 | 0.051 | 0.103 | 0.069 | 0.077 | 0.057 | 0.072 | 0.054 |
| L External capsule | 0.079 | 0.052 | 0.062 | 0.052 | 0.115 | 0.077 | 0.075 | 0.073 | 0.100 | 0.071 | 0.040 | 0.047 | 0.062 | 0.061 | 0.106 | 0.066 | 0.064 | 0.062 | 0.068 | 0.065 |
| R Cingulum (cingulate gyrus) | 0.044 | 0.050 | 0.035 | 0.043 | 0.065 | 0.051 | 0.037 | 0.034 | 0.078 | 0.046 | 0.029 | 0.035 | 0.043 | 0.044 | 0.091 | 0.068 | 0.094 | 0.056 | 0.063 | 0.065 |
| L Cingulum (cingulate gyrus) | 0.058 | 0.048 | 0.056 | 0.039 | 0.059 | 0.057 | 0.037 | 0.041 | 0.085 | 0.039 | 0.029 | 0.030 | 0.051 | 0.037 | 0.088 | 0.064 | 0.074 | 0.054 | 0.060 | 0.060 |
| R Cingulum (hippocampus) | 0.066 | 0.054 | 0.049 | 0.043 | 0.093 | 0.075 | 0.045 | 0.084 | 0.040 | 0.057 | 0.016 | 0.042 | 0.018 | 0.036 | 0.101 | 0.051 | 0.067 | 0.053 | 0.075 | 0.047 |
| L Cingulum (hippocampus) | 0.078 | 0.082 | 0.074 | 0.069 | 0.090 | 0.077 | 0.061 | 0.085 | 0.042 | 0.059 | 0.029 | 0.059 | 0.043 | 0.068 | 0.060 | 0.052 | 0.053 | 0.046 | 0.032 | 0.040 |
| R Fornix / Stria terminalis | 0.093 | 0.042 | 0.081 | 0.055 | 0.123 | 0.067 | 0.048 | 0.046 | 0.081 | 0.048 | 0.046 | 0.043 | 0.050 | 0.051 | 0.183 | 0.040 | 0.100 | 0.044 | 0.113 | 0.040 |
| L Fornix / Stria terminalis | 0.096 | 0.051 | 0.083 | 0.053 | 0.106 | 0.066 | 0.082 | 0.057 | 0.126 | 0.070 | 0.061 | 0.062 | 0.097 | 0.076 | 0.167 | 0.052 | 0.095 | 0.043 | 0.106 | 0.039 |
| R Superior longitudinal fasciculus | 0.132 | 0.045 | 0.104 | 0.035 | 0.115 | 0.048 | 0.092 | 0.045 | 0.110 | 0.039 | 0.083 | 0.045 | 0.082 | 0.042 | 0.137 | 0.036 | 0.146 | 0.042 | 0.147 | 0.034 |
| L Superior longitudinal fasciculus | 0.108 | 0.043 | 0.101 | 0.047 | 0.099 | 0.032 | 0.092 | 0.030 | 0.109 | 0.028 | 0.068 | 0.026 | 0.093 | 0.026 | 0.140 | 0.041 | 0.120 | 0.038 | 0.123 | 0.040 |
| R Superior fronto-occipital fasciculus | 0.091 | 0.025 | 0.076 | 0.031 | 0.102 | 0.027 | 0.081 | 0.032 | 0.071 | 0.034 | 0.035 | 0.028 | 0.044 | 0.035 | 0.087 | 0.044 | 0.074 | 0.031 | 0.077 | 0.022 |
| L Superior fronto-occipital fasciculus | 0.088 | 0.026 | 0.092 | 0.035 | 0.043 | 0.038 | 0.040 | 0.033 | 0.106 | 0.035 | 0.052 | 0.036 | 0.080 | 0.029 | 0.086 | 0.039 | 0.053 | 0.035 | 0.056 | 0.030 |
| R Inferior fronto-occipital fasciculus | 0.073 | 0.041 | 0.087 | 0.053 | 0.088 | 0.059 | 0.034 | 0.046 | 0.054 | 0.043 | 0.024 | 0.032 | 0.020 | 0.036 | 0.078 | 0.057 | 0.070 | 0.048 | 0.070 | 0.059 |
| L Inferior fronto-occipital fasciculus | 0.066 | 0.038 | 0.060 | 0.045 | 0.084 | 0.065 | 0.053 | 0.045 | 0.059 | 0.046 | 0.013 | 0.026 | 0.021 | 0.037 | 0.074 | 0.049 | 0.027 | 0.040 | 0.043 | 0.048 |
| R Uncinate fasciculus | 0.047 | 0.034 | 0.045 | 0.035 | 0.052 | 0.041 | 0.009 | 0.015 | 0.006 | 0.016 | 0.001 | 0.004 | 0.007 | 0.017 | 0.070 | 0.037 | 0.053 | 0.031 | 0.041 | 0.029 |
| L Uncinate fasciculus | 0.040 | 0.025 | 0.039 | 0.031 | 0.028 | 0.036 | 0.008 | 0.017 | 0.004 | 0.010 | 0.000 | 0.000 | 0.000 | 0.000 | 0.032 | 0.029 | 0.000 | 0.002 | 0.006 | 0.014 |
| R Tapetum | 0.028 | 0.023 | 0.019 | 0.024 | 0.006 | 0.011 | 0.004 | 0.008 | 0.041 | 0.027 | 0.007 | 0.017 | 0.030 | 0.026 | 0.054 | 0.030 | 0.048 | 0.030 | 0.012 | 0.021 |

**Table S3.** Runners individual data.

|  | **Runner 1** | **Runner 2** | **Runner 3** | **Runner 4** |
| --- | --- | --- | --- | --- |
| **Age** | 73 | 59 | 45 | 46 |
| **Weight (kg)** | 58 | 76 | 69 | 71 |
| **Height (cm)** | 160 | 181 | 180 | 180 |
| **Time to finish line** | 3h 45m | 3h 59m | 4h 40m | 6h 06m |
| **Heart rate (average bpm)** | 150 | 145 | 152 | Not assessed |
| **Ambient Temperature (°C)** | 17-24 | 17-24 | 13-24 | 13-24 |
| **Estimated KCalories burned** | 2393 | 3364 | 2776 | Not assessed |

## Notes

### Competing Interest Statement

The authors have declared no competing interest.

